# Connexin 43 plays an important role in the transformation of human cholangiocytes upon stimulation with *Clonochis sinensis* excretory-secretory protein and *N*-nitrosodimethylamine

**DOI:** 10.1101/418350

**Authors:** Eun-Min Kim, Young Mee Bae, Min-Ho Choi, Sung-Tae Hong

## Abstract

**Background:** *Clonorchis sinensis* is a group I bio-carcinogen responsible for cholangiocarcinoma (CHCA) in humans. However, the mechanism by which *C. sinensis* promotes carcinogenesis is unclear.

**Methodology:** Using the human cholangiocyte line H69, we investigated cell proliferation and gap junction protein expression after stimulation with the hepatotoxin *N*-nitrosodimethylamine (NDMA) and/or excretory-secretory products (ESP) of C. *sinensis*, which induce inflammation. NDMA and ESP treatment increased proliferation by 146% and the proportion of cells in the G2/M phase by 37%. Moreover, the expression of the cell cycle protein E2F1 and the cell proliferation-related proteins Ki-67 and cytokeratin 19 increased in response to combined treatment with NDMA and ESP. The gap-junction proteins connexin (Cx) 43 and Cx26 also increased. In contrast, Cx32 expression decreased in cells treated with NDMA and ESP. Cox-2 was also upregulated. Silencing of Cx43 reduced cell proliferation and significantly suppressed Cx26 and Cox-2 expression.

**Conclusions:** These results suggest that Cx43 is an important factor in CHCA induced by *C. sinensis* ESP and NDMA and further investigations targeting this pathway may allow prevention of this deadly disease.

**Author summary:** *Clonorchis sinensis*, a human fluke, resides in the liver of humans and is commonly found in the common bile duct and gall bladder. This parasite is the main cause of cholangiocarcinoma, also called bile duct cancer, in humans. Of note, the excretory-secretory products (ESP) of *C. sinensis* are known to cause inflammation in the biliary epithelium, which may ultimately result in neoplasms via production of reactive oxygen species and subsequent DNA damage. Together with *N*-nitrosodimethylamine (NDMA), a potent hepatotoxin that can cause fibrosis and tumors in the liver, ESP led to an increase in the growth and proliferation of cholangiocytes. Our results showed that the ESPs of *C. sinensis* induced pro-inflammatory responses by increasing the levels of proinflammatory cytokines and nuclear factor kappa B (NFκB), which in turn, enhanced the production of connexin 43 (Cx43), a gap-junction protein. Therefore, Cx 43 can serve as a potential target for developing a therapeutic strategy for the treatment of cholangiocarcinoma in humans.

## Introduction

*Clonorchis sinensis* is a human liver fluke that induces cholangiocarcinoma (CHCA) in humans [1]. Clonorchiasis has been endemic in Asia for a long time, especially among residents who live along rivers and consume raw freshwater fish [2].

The mechanism by which *C. sinensis* induces CHCA is not well understood, but chronic hepatobiliary damage, a precursor to CHCA, in clonorchiasis is a multi-factorial outcome of the mechanical and biochemical irritation of the biliary epithelium by flukes via their suckers, metabolites, and excretory-secretory products (ESP) [3]. Local inflammation and the systemic immune response in the host [4-7] produce reactive oxygen species and reactive nitrogen compounds, which may cause DNA damage, leading to neoplasms [8, 9].

*N*-nitrosodimethylamine (NDMA) is a potent hepatotoxin that can cause fibrosis and tumors in the liver of rats via the activation of CYP450 enzymes [10] and hamsters infected with *C. sinensis* are at a greater risk of developing NDMA-induced or inflammation-mediated CHCA than uninfected hamsters [11, 12]. Previously, we reported that exposure to NDMA and the ESP of *C. sinensis* increases HEK293T cell proliferation and the proportion of cells in the G2/M phase [3].

CHCA is potentially caused by increased levels of proinflammatory cytokines and nuclear factor kappa B (NFκB), which regulate the activities of cyclooxygenase-2 (Cox-2) and inducible nitric oxide synthase, and disturb the homeostasis of oxidants/anti-oxidants and DNA repair enzymes [13]. ESPs of *C. sinensis* induce pro-inflammatory responses *in vitro* by the upregulation of TLR4 and its downstream transduction signals, including MyD88-dependent IκB-α degradation and NFκB activation [14, 15]. NFκB may also influence the production of Cx43, a gap-junction protein, in liver cirrhosis [16, 17].

Gap junctions are clusters of transmembrane channels on the cell membrane that permit direct intercellular exchange of ions, secondary messengers, and small signaling molecules influencing cell growth, differentiation, and cancerous changes [17-21]. Among gap junction proteins, Cx43 is involved in almost all steps of the inflammatory response of cells, cytokine production, and inflammatory cell migration [17, 20, 22]. The substantial role of Cx43 in carcinogenesis is highlighted by the fact that high levels of Cx-43 expression increase the invasion of breast tumor cells and promote tumors in melanoma [22].

Alterations of Cx expression have been reported in cancer [21, 23]. In hepatocellular carcinoma (HCC), for instance, reduced Cx32 expression is accompanied by increased expression of Cx43, which promotes HCC via cell-to-cell communication [16, 20, 23]. Fujimoto et al. [23] have shown that Cx32 has a suppressive effect in metastatic renal cell carcinoma. However, the role of connexins in cancer is still controversial [23], and the influence of gap junctions in CHCA caused by *C. sinensis* has not yet been examined.

In this study, to determine the mechanisms underlying the carcinogenic effects of ESP from *C. sinensis,* we investigated changes in cell proliferation, proinflammatory molecules, and connexin production in cholangiocytes (H69 cell line) exposed to ESP from *C. sinensis* and the carcinogen NDMA. We found that the silencing of Cx43 reduced ESP- and NDMA-induced cell proliferation and the expression of Cox-2.

## Methods

### Preparation of ESP

#### Animals

The animal experimental protocol was approved and reviewed by the Institutional Animal Care and Use Committee (IACUC) of Seoul National University Health System, Seoul, Korea (approval no. SNU-060511-1) and followed the National Institutes of Health (NIH) guideline for the care and use of laboratory animals (NIH publication no. 85–23, 1985, revised 1996). The facility is accredited by the Ministry of Food and Drug Administration and by the Ministry of Education, Science and Technology (LNL08-402) as an animal experiment facility. Male Sprague–Dawley rats at 6 weeks of age were purchased from Koatech Co. (Seoul, Korea) and housed in an ABL-2 animal facility at Seoul National University College of Medicine. All rats were bred in filter cages under positive pressure according to institutionally approved guidelines.

#### Recovery of metacercariae of *C. sinensis*

*Pseudorasbora parva*, the second intermediate host of C. sinensis, which was naturally infected *with C. sinensis*, was purchased from Sancheong-*gun, Gyeong*sangnam-do, Republic of Korea, an endemic area for clonorchiasis. Metacercariae of *C. sinensis* were collected after the diges*tion of fis*h with pepsin-HCl (0.6%) artificial gastric juice for 1 h at 37°C.

#### Infection of experimental animals with *C. sinensis* and collection of ESP

Sprague–Dawley rats were individually infected orally with 50 metacercariae of *C. sinensis*. Eight weeks post-infection, adult worms were collected from b*ile ducts a*nd washed several times with phosphate-buffered saline (PBS). The freshly isolated worms were then incubated in sterile PBS containing antibiotics for 24 h in an atmosphere of 5% CO_2_ at 37°C. After incubation, the medium was centrifuged for 10 min at 800 _r_pm to remove the worms and debris. The supernatant was then further centrifuged for 10 min at 3000 rpm and filtered with a syringe-driven 0.45-µm filter unit. The amount of protein in each extract was measured using the Bradford assay (Thermo, Rockford, IL, USA). The concentration of endotoxin (LPS) was measured using an LAL QCL-1000 Kit (LONZA, Switzerland), and as a result, the LPS contained in 10 µg/mL of ESP was measured to be less than 0.001 (EU). Therefore, there is no effect of LPS on the results of this paper.

### Cell culture and experimental design

#### Cell culture

SV40-transformed human cholangiocytes (H69) from Dr. Dae-Gon Kim of Chonbuk National University for providing were divided into four treatment conditions and cultured for more than 180 days; the medium was replaced every 72 h. The cells were then treated as follows: control, cultured in plain medium; 100 ng/mL NDMA, cultured in medium containing 100 ng/mL NDMA; ESP, cultured in medium containing 10 µg/mL ESP; and NDMA + ESP, cultured in medium containing 10 µg/mL NDMA and 100 ng/mL ESP. H69 cells were cultured in Dulbecco’s modified Eagle’s medium (DMEM; Gibco, Carlsbad, CA, USA) and DMEM/F12 supplemented with 10% heat-inactivated fetal bovine serum (FBS; Gibco), 2 mM L-glutamine, 100 µg/mL penicillin, 0.243 mg/mL adenine (Sigma, St. Louis, MO, USA; A68626), 5 µg/mL insulin (Sigma; I6634), 10 µg/mL epinephrine (Sigma; E4250), 5 µg/mL Triiodonine-transferrin (Sigma; T8158), 30 ng/mL epidermal growth factor (R&D Systems, Minneapolis, MN, USA; 236-EG), 1.1 µM hydrocortisone, and 100 U/mL streptomycin at 37°C in a humidified atmosphere of 5% CO_2_.

#### Cell proliferation assay

The PrestoBlue cell viability reagent was utilized to evaluate cell proliferation. For each assay, cells were seeded at a density of 5 × 10^3^ cells/well on 96-well plates. After 24 h of incubation, the medium was replaced with 2% FBS-DMEM without phenol red. The cells were then incubated in the presence of PBS (vehicle) or with 100 ng/mL NDMA with or without 10 µg/mL ESP for another 72 h. The PrestoBlue cell viability reagent (1 mg/mL) was dissolved in warm medium, and 1.25 mM phenazine methosulfate (PMS) was prepared in PBS. Following incubation for the indicated periods, 50 µL of the PrestoBlue cell viability reagent was added to each well. The plates were then incubated for 1 h. The conversion of PrestoBlue cell viability reagent was quantified by measuring the absorbance at 570 and 600 nm using a microtiter plate reader.

#### Cell cycle analysis

For the cell cycle analysis, H69 cells were plated in six-well culture plates at 2 × 10^5^ cells/well in 2 mL of DMEM containing 10% FBS. They were then treated with 100 ng/mL NDMA with or without 10 µg/mL ESP for 72 h and stained with propidium iodide (PI). The PI-stained cells were analyzed using a FACSCalibur multicolor flow cytometer (Becton-Dickinson, Franklin lakes, NJ, USA), and data were analyzed using CellQuest software (Becton-Dickinson).

#### Western blotting

For western blots, cells were lysed using 1% Nonidet P-40 in a buffer containing 150 mM NaCl, 10 mM NaF, 1 mM PMSF, 200 µM Na_3_VO_4_, and 50 mM HEPES, pH 7.4. Equal amounts of protein were separated by 8% and 10% sodium dodecyl sulfate-polyacrylamide gel electrophoresis (SDS-PAGE) and transferred to polyvinylidene fluoride (PVDF) membranes (Immobilon; Millipore, Billerica, MA, USA). The membranes were then probed with antibodies against E2F1, Ki-67, Ck19, Cox-2, connexin 43, connexin 32, connexin 26, and calnexin. The primary antibodies were detected using goat anti-rabbit or rabbit anti-mouse secondary antibodies conjugated with HRP and visualized using an enhanced chemiluminescence kit (ECL; Amersham Pharmacia Biotech, North Massapequa, NY, USA). The western blotting results were obtained by a densitometric analysis using ImageJ (NIH, Bethesda, MD, USA).

#### Antibodies

Polyclonal or monoclonal antibodies were used to detect the expression of cell-cycle-related proteins, including anti-E2F1 (sc-193; Santa Cruz Biotechnology, Santa Cruz, CA, USA) and anti-Ki-67 (SP6; Abcam, Cambridge, MA, USA). The other antibodies included anti-Cox-2 (c-9897; Cayman Chemicals, Ann Arbor, MI, USA) and anti-cytokeratin-19 (Ab53119; Abcam) as cancer-related makers, and anti-connexin 26 (138100; Invitrogen, Carlsbad, CA, USA), anti-connexin 32 (358900; Invitrogen), and anti-connexin 43 (138300; Invitrogen). An antibody against calnexin (BD 610523) used as a control was purchased from Transduction Laboratories (BD Biosciences, San Jose, CA, USA) and used at a 1:1,000 dilution. Anti-mouse, anti-rabbit, and anti-goat IgG antisera conjugated with horseradish peroxidase (HRP) were purchased from DAKO (Glostrup, Denmark).

#### Confocal microscopy

Cells were washed with cold PBS three times and fixed with 2% paraformaldehyde in PBS for 30 min. Permeabilization was performed by treating the cells with 0.2% (w/v) Triton X-100 in PBS for 5 min and then blocking with 0.5% BSA in PBS for 1 h. After blocking, the cells were incubated with primary antibodies against connexin 26, 32, and 43 (Invitrogen) diluted in BSA-PBS at 25°C for 2 h and then incubated in secondary antibodies diluted in BSA-PBS at room temperature for 30 min. After washing with 1× PBS, the cells were stained with DAPI and observed under a confocal laser scanning microscope (LSM PASCAL; Carl Zeiss, Jena, Germany).

### Cx43 silencing with siRNA

Three selected human Cx43-siRNAs (TriFECTa Kit disRNA Duplex, IDT, San Jose, CA, USA), with negative and positive controls, and specific siRNA targeting connexin 43 (NM-00165) (Table 1) were purchased from IDT. The transfection experiments were performed using the TransIT-TKO Kit (Mirus, Madison, WI, USA) following the manufacturer’s instructions. Briefly, a 25 nM siRNA solution was mixed with 10 µL of TransIT-TKO and added to the wells of a 6-well plate containing 2 × 10^5^ H69 cells for 72 h. Cells were then treated with NDMA, ESP of *C. sinensis*, or the combination for 72 h. Medium and cells (rinsed 2 times with ice-cold PBS) were harvested 3 days later. The efficiency of transfection was assessed by measuring the expression of gap junction proteins (connexin 26, connexin 32, and connexin 43) by real-time PCR.

**Table 1.**
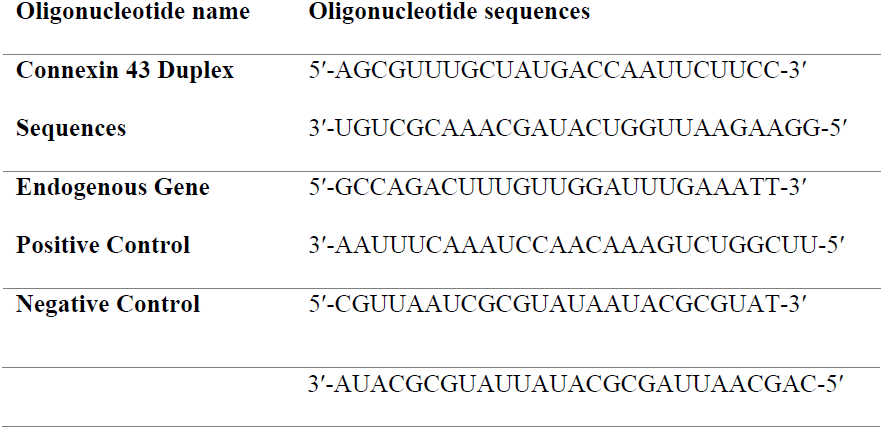
Connexin43-specific siRNA oligonucleotide sequences

### Real-time PCR

RNA samples from each cell line were column-purified using the RNeasy Mini Kit (Qiagen, Hilden, Germany). Reverse transcription and real-time PCR (RT-PCR) were performed to determine the mRNA expression levels of *Cx26, Cx32, Cx43*, and *Cox-2*, using *GAPDH* as a control (Applied Biosystems, Santa Clara, CA, USA). The thermal cycling parameters for reverse transcription were modified according to the Applied Biosystems manual. Hexamer incubation at 25°C for 10 min and reverse transcription at 42°C for 30 min was followed by reverse transcriptase inactivation at 95°C for 5 min. cDNA (20 ng) from the previous step was subjected to RT-PCR using specific sets of primers (Table 2 in a total reaction volume of 25 µL (Applied Biosystems). RT-PCR was performed in an optical 96-well plate using an ABI PRISM 7900 HT Sequence Detection System (Applied Biosystems) and TaqMan probe detection chemistry. The running protocol was as follows: initial denaturation at 95°C for 10 min, and 40 cycles of amplification at 95°C for 15 s and 60°C for 1 min. After PCR, a dissociation curve was constructed by increasing the temperature from 65°C to 95°C to evaluate the PCR amplification specificity. The cycle threshold (Ct) value was recorded for each sample.

**Table 2.**
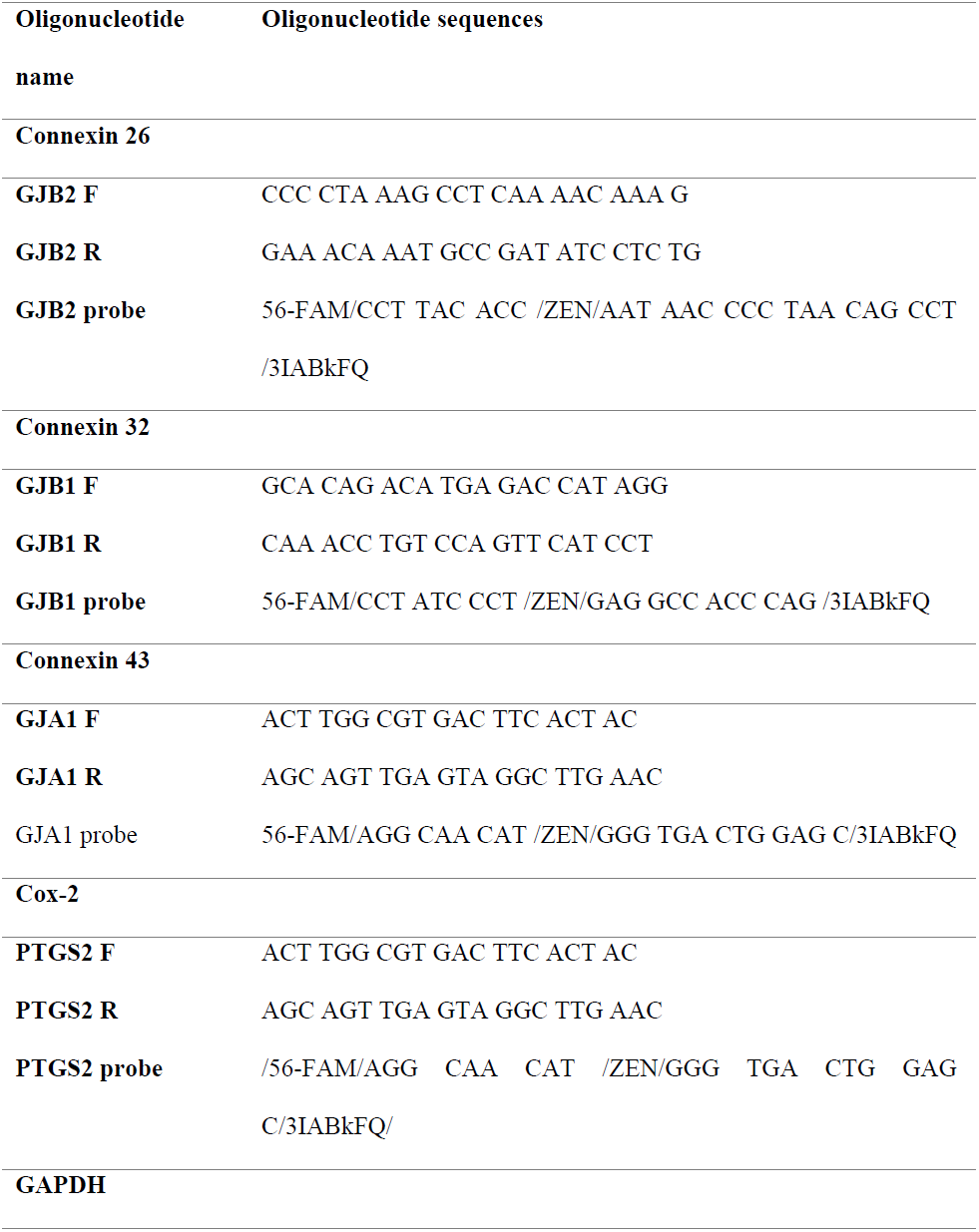
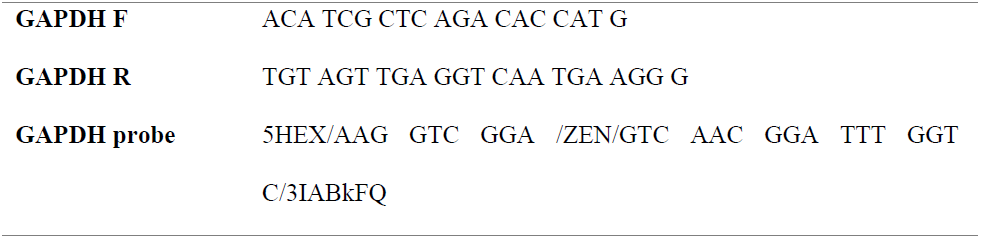
Oligonucleotide primers and detection probe for real-time PCR

### Statistical analysis

Statistical significance was analyzed using Student’s *t*-tests. Differences were considered statistically significant at a ^*^*P* < 0.05, ^**^*P* < 0.01 and ^***^*P* <0.001 versus control. Data are presented as the mean ± standard error of the mean (SEM) from at least three independent experiments.

## Results

### ESP and NDMA synergistically increase H69 cell proliferation

The roles of NDMA and ESP in the proliferation of H69 cells, as determined by cell viability, were investigated. Proliferation was higher in H69 cells treated with NDMA and ESP of *C. sinensis* than in control cells. Compared to control cells, the average increase for cells treated with NDMA was 112%, for cells treated with ESP was 120% (*P* < 0.05), and for cells treated with NDMA + ESP was 146% (*P* < 0.001). NDMA and ESP had synergistic effects on cell proliferation (Fig 1A).

**Fig 1:**
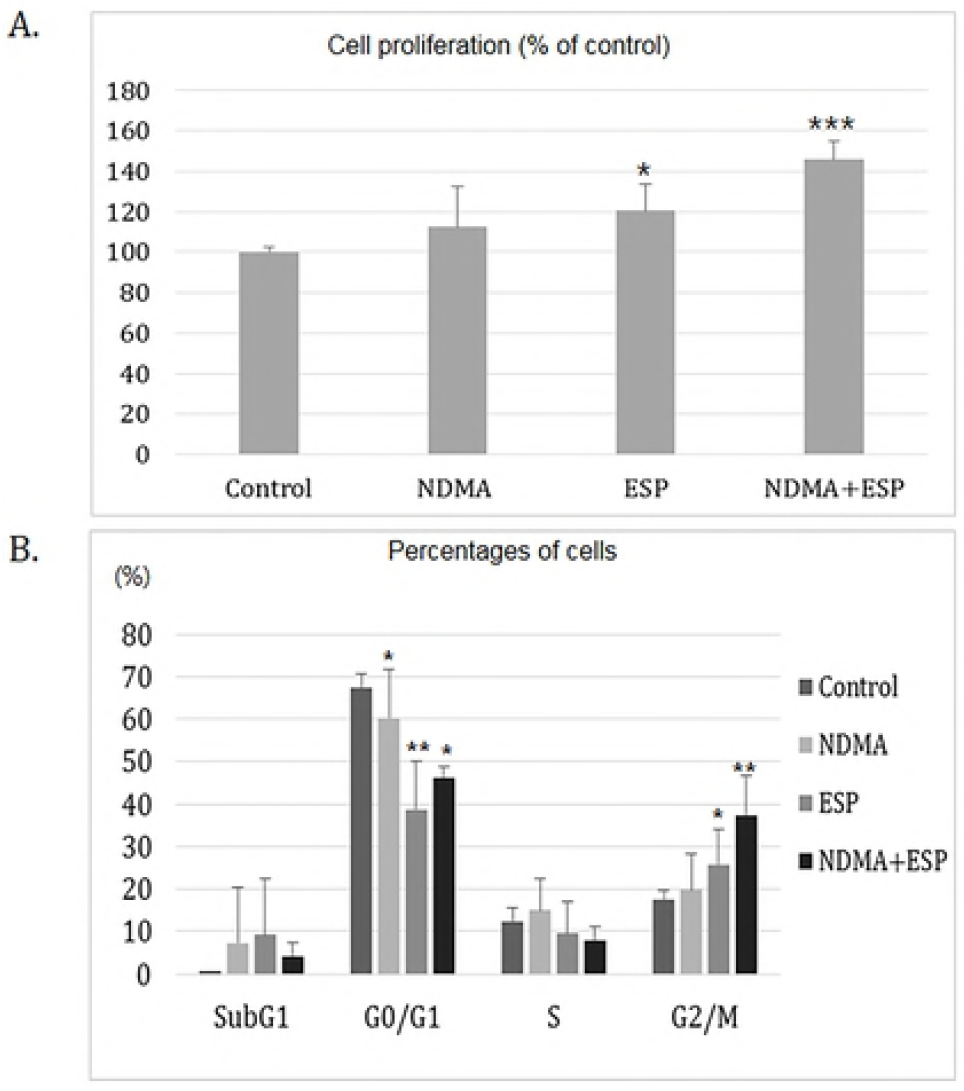
Effects of NDMA and ESP of *C. sinensis* on cell proliferation and cell cycle progression in human cholangiocytes (H69 cells). **A.** Cells were plated in 96-well plates (5 × 10^3^ cells/well). Cell proliferation for each treatment was determined using the PrestoBlue cell viability assay. **B.** After H69 cells were treated with NDMA and ESP for 72 h, PI staining was performed to determine the percentage of cells in each phase. Data represent the mean ± SE of five independent experiments. ^*^*P* < 0.05, ^**^*P* < 0.01 and ^***^*P* < 0.001 versus control.

### Cell cycle distribution upon NDMA and/or ESP treatment in H69 cells

Cell cycle progression was monitored using propidium iodide (PI) staining (Fig 1B). In H69 cells treated with NDMA, ESP, or NDMA + ESP for 72 h, the number of cells in the G0/G1 phase significantly decreased, but cells in the G2/M phase increased significantly compared to cell counts in the control group (Fig 1B). Fewer S-phase cells were identified in the ESP- and NDMA + ESP-treated cells than in control cells. There is no significant increase in G2/M phase in cells treated with NDMA compared to control group (Fig 1B). The proportions of G2/M-phase cells in each condition were as follows: control, 17%; NDMA, 20%; ESP, 26%; NDMA + ESP, 37% (*P* = 0.007).

### Upregulation of inflammation- or transformation-related proteins by NDMA and ESP in H69 cells

Immunoblotting was utilized to detect the regulation of cell-cycle-related proteins in each group using calnexin as a loading control. The expression levels of several cell proliferation- and inflammation-related proteins, including E2F1, Ki-67, Cy-19, and Cox-2 (an essential regulator of the G2/M transition), were upregulated, especially in NDMA + ESP-treated cells (Fig 2).

**Fig 2:**
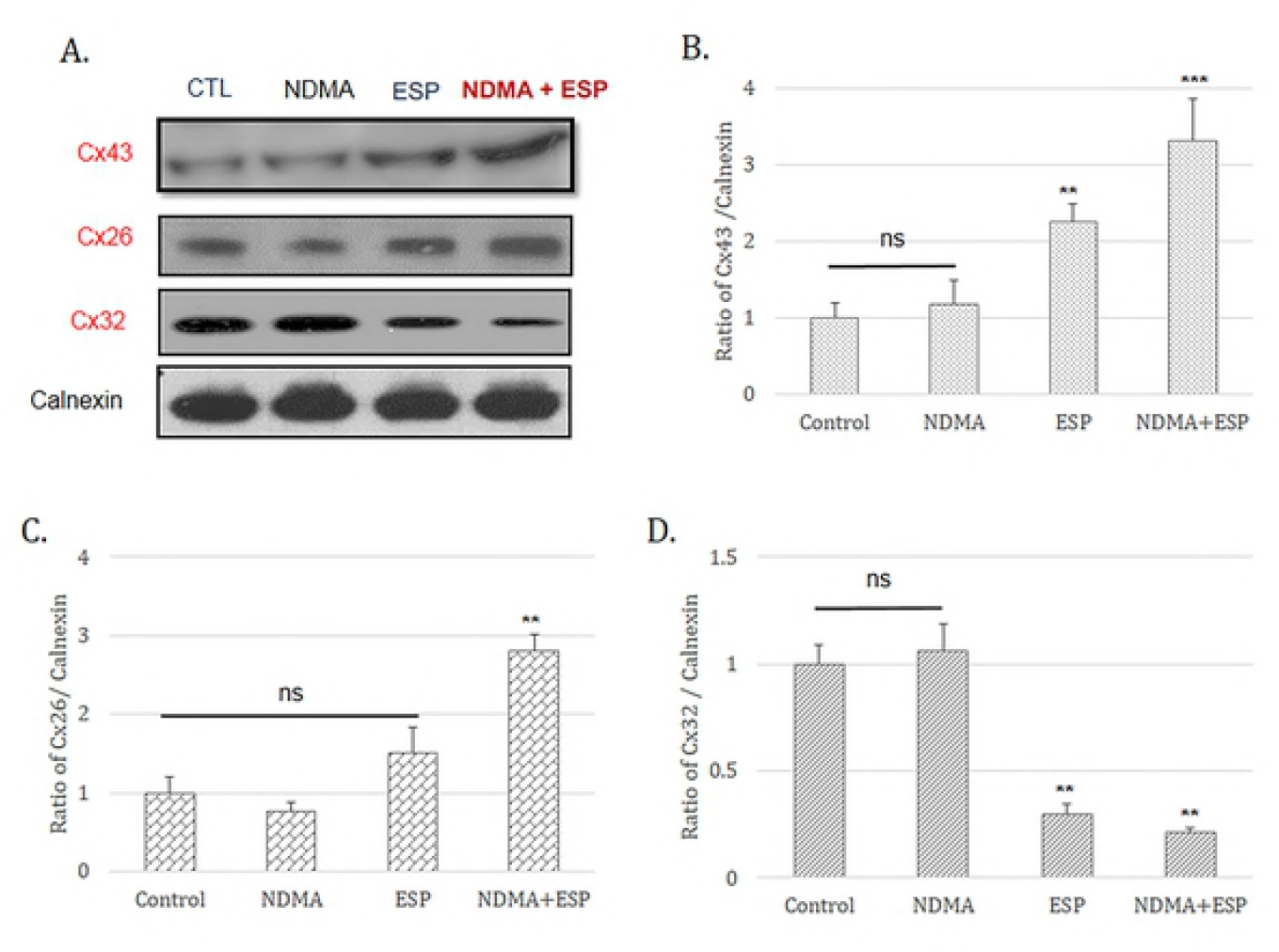
Expression of inflammation- or transformation-related proteins in H69 cells after treatment with NDMA and ESP, assessed by western blotting. H69 cells were incubated with PBS (vehicle), NDMA, ESP, or both for 72 h. The cells were collected for protein extraction. The blots of each groups were run under same experimental conditions and the images were cropped from different parts of the same gels, DATA represent the mean ± SE of five independent experiments.^*\**^*P* < 0.05, ^*\*\**^*P* < 0.01 and ^*\*\*\**^*P* < 0.001 versus control.

### Gap junction proteins in H69 cells

The expression levels of Cx26 and Cx43 were high in NDMA + ESP-treated cells, as confirmed by immunoblotting (Fig 3). The intracellular concentrations of Cx26 and Cx43 in each condition were observed under a confocal microscope. Western blotting indicated increases in Cx26 and Cx43 expression in cells treated with ESP and NDMA + ESP (Figs 3). The expression of Cx32 (blue) was markedly lower in cells treated with NDMA + ESP than in other cells. Immunofluorescence confocal microscopy indicated that the intracellular concentrations of Cx26 (green) and Cx43 (green) increased in cells treated with NDMA + ESP (Fig 4).

**Fig 3:**
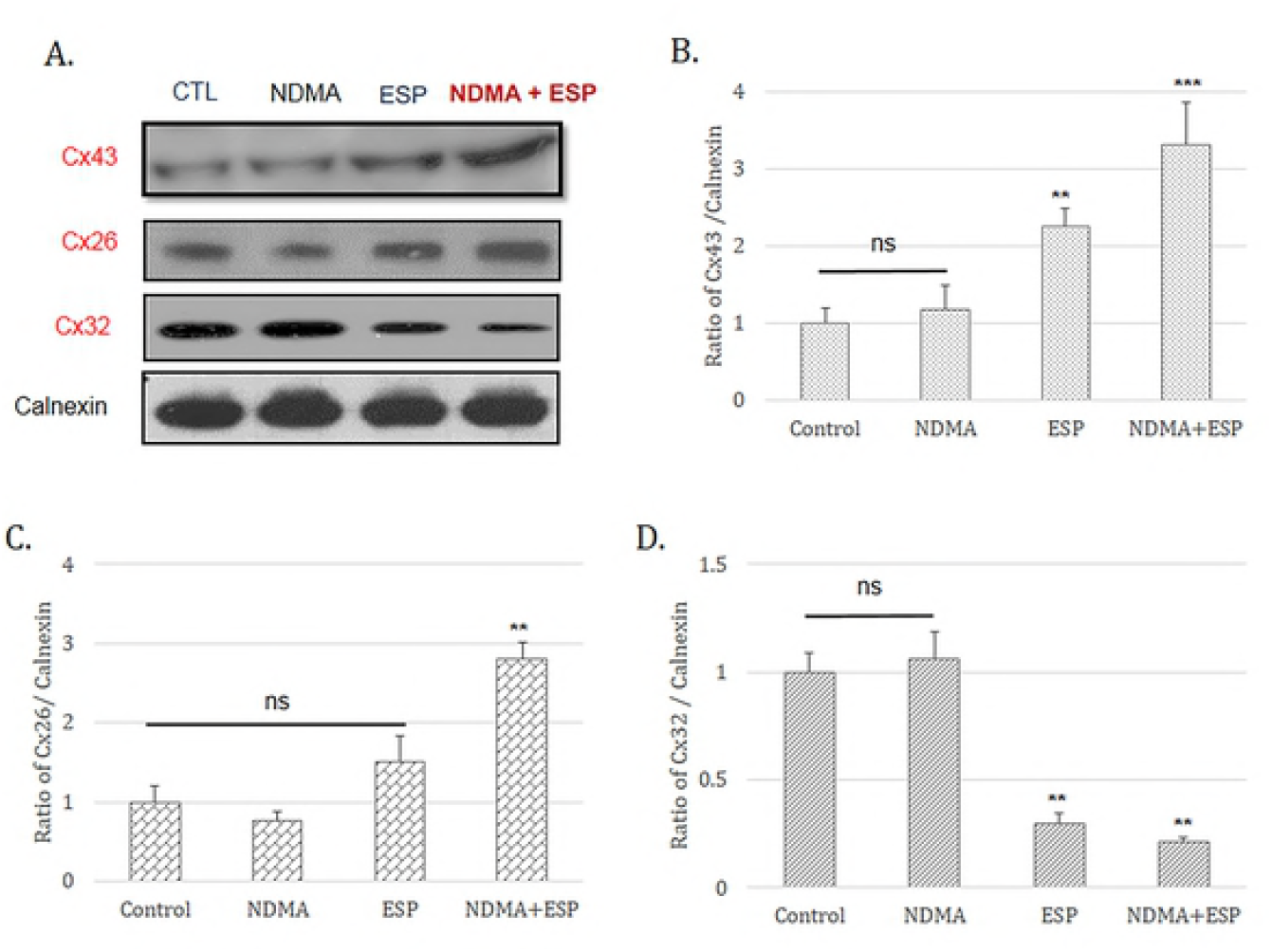
Expression of the gap-junction proteins connexin 26, connexin 32, and connexin 43 in H69 cells after treatment with NDMA and/or ESP, as determined by western blotting. H69 cells were incubated with either PBS (vehicle) or NDMA and/or ESP for 72 h, and the cells were collected for protein extraction. The blots of each groups were run under same experimental conditions and the images were cropped from different parts of the same gels, DATA represent the mean ± SE of five independent experiments. ^*^*P* < 0.05, ^**^*P* < 0.01 and ^***^*P* < 0.001 versus control.

**Fig 4:**
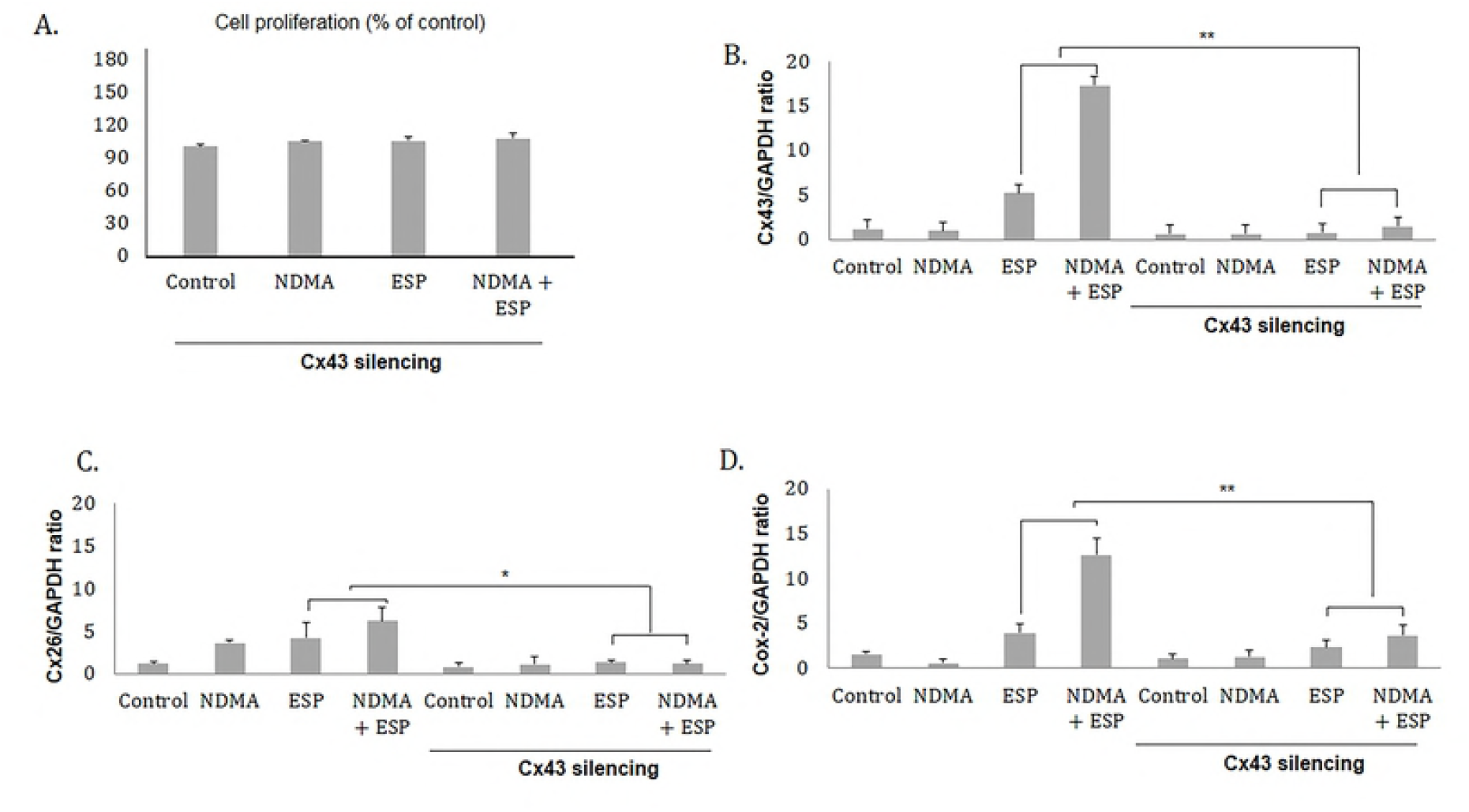
Concentration of intracellular connexin 26, connexin 32, and connexin 43 in human cholangiocytes (H69 cells). After treatment with NDMA and ESP for 72 h, the concentration of intracellular connexin 26 in H69 cells was measured by laser scanning microscopy (×4,000). Scale bar = 25 µm. Data represent the mean ± SE of five independent experiments. ^*^*P* < 0.05, ^**^*P* < 0.01 and ^***^*P* < 0.001 versus control.

### Reduced cell proliferation upon NDMA and ESP treatment in H69 cells by Cx43 silencing compare to silencing negative control

Downregulation of Cx43 by Cx43-specific small interfering RNA resulted in all groups were not significant, even though NDMA and ESP stimulated cells. The average increases compared to the control were as follows: NDMA, 104.5%; ESP, 105.1%; and NDMA + ESP, 107.6% (Fig 5A).

**Fig 5:**
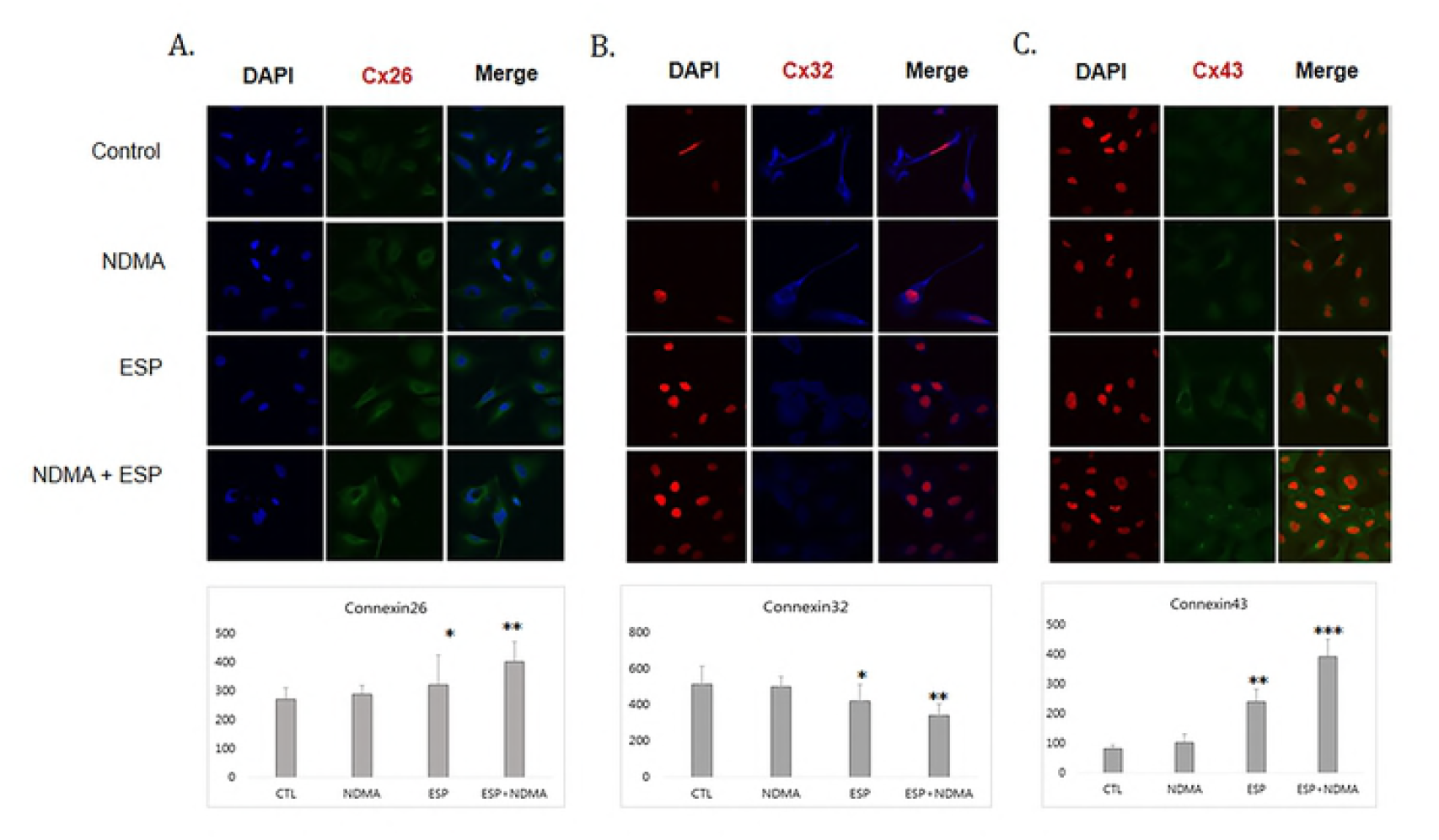
Effect of connexin 43 silencing in H69 cells. **A.** Reduced cell proliferation upon NDMA and ESP treatment in H69 cells by connexin 43 silencing. **B.** Uptake of Cx43 siRNA reduces *Cx43* expression, as confirmed by real-time PCR. *Cx43* expression was remarkably reduced in H69 cells transfected with *Cx43*-specific siRNA. **C.** Ratio of *Cx26/GAPDH* in H69 cells transfected with *Cx43*-specific siRNA. *Cx26* expression was remarkably reduced in H69 cells transfected with *Cx43*-specific siRNA. **D.** The ratio of *Cox-2/GAPDH* in H69 cells transfected with *Cx43*-specific siRNA. *Cox-2* expression was remarkably reduced in H69 cells transfected with *Cx43*-specific siRNA. Data represent the mean ± SE of five independent experiments. ^*^*P* < 0.001 versus control siRNA.

### Downregulation of Cx43 by Cx43-specific small interfering RNA reduced the expression of Cx26 and Cox-2 in H69 cells

To evaluate the effect of Cx43 downregulation on the expression of other gap junction proteins and Cox-2, H69 cells were harvested after treatment with NDMA and ESP of *C. sinensis* for 72 h. Transfection with Cx43 siRNA resulted in a reduction in *Cx43* expression of greater than 70% when compared with that of the liposome-only control (Fig 5A). Real-time PCR showed that Cx43 silencing resulted in the downregulation of *Cx26* and *Cox-2* (Figs 5C and 5D, respectively). However, *Cx32* expression was not affected (not shown).

## Discussion

Our results provided the first evidence for the involvement of gap junction proteins in the pathogenesis of CHCA by *C. sinensis.* NDMA and ESP of *C. sinensis* increased Cx43 expression substantially in H69 human cholangiocytes. In addition, in the presence of ESP and NDMA stimulation, *Cx43* knockdown inhibited *Cox-2* expression in H69 cells. Yan et al. [15] reported that iNOS is highly expressed in Kupffer cells, sinusoidal endothelial cells, and biliary epithelial cells in BALB/c mice after *C. sinensis* infection. Nitric oxide (NO) formation and nitrosation may contribute to the development of *C. sinensis*-associated carcinogenesis [11, 12, 14, 15]. The induction of iNOS under inflammatory conditions suggests that NO is involved in the upregulation of *Cx43* [16, 19]. Therefore, we also expect the involvement of iNOS in the elevation of *Cx43* expression under inflammatory conditions in the present *C sinensis* model. Further studies are warranted to test this hypothesis.

Our results indicated that ESP of *C. sinensis* and NDMA had a synergistic effect on the proliferation of human cholangiocytes (Fig 1). In addition to the observed increase in cell proliferation and alteration of the cell cycle, the expression of the gap junction proteins Cx43 and Cx26 increased in H69 cells treated with NDMA and ESP. Most normal cells have functional gap junctional intercellular communication, in contrast to the dysfunctional communication of cancer cells [17-22]. When used in combination with NDMA, ESP of *C. sinensis* induced cell proliferation and increased the expression of E2F1, Ck19, and Ki-67. When H69 cells were co-stimulated with NDMA and ESP, cell proliferation increased. Additionally, as shown in Fig 1B, treatment with NDMA + ESP maximized the proportion of cells in the G2/M phase, implying that NDMA and ESP synergistically stimulate cell cycle progression. We analyzed the expression of a number of cell proliferation-, cell cycle-, and inflammation-related proteins (Fig 2), including E2F1, Ki-67, and Cy19 [24-26]. E2F1 is able to induce cell cycle progression, resulting in cell proliferation [3, 24]. Consistent with these previous findings, we observed that increased expression of E2F1 stimulated cell proliferation. C-Met is involved in early events of carcinogenesis, and Ki-67 is involved in the formation of invasive carcinoma [25-28]. Biliary epithelial cells retain Cy19 expression after neoplastic transformation in almost all cases [26, 28]. In our study, Cy19 and Ki-67 were upregulated in response to the stimulation of H69 cell proliferation.

Cyclooxygenase 2 (Cox-2), an enzyme involved in the production of prostaglandins, was over-expressed when cells were stimulated by NDMA and ESP. Cox-2 over-expression has been observed in various inflammatory diseases and in bile duct carcinoma cells, mainly in the cytoplasm [29-30]. Importantly, bile duct epithelial cells in primary sclerosing cholangitis show very strong Cox-2 expression, comparable to that in carcinoma cells. In contrast, epithelial cells in primary biliary cirrhosis show moderate levels of Cox-2 expression [29, 30]. In this context, the over-expression of Cy19, Ki-67, and Cox-2 may result in the transformation of normal H69 cells to cancer-like cells by stimulation with NDMA and ESP of *C. sinensis*. In the present study, Cx43 and Cx26 expression levels were increased in H69 cells upon stimulation with NDMA and ESP of *C. sinensis*. In contrast, Cx32 was significantly downregulated. Increased expression of hepatic Cx43 was noted in cirrhosis and in a mouse model of acute-on-chronic liver failure in response to LPS, and this effect was related to the severity of inflammation [19]. This increased Cx43 expression was likely an adaptive protective response of the liver to enable better cell-to-cell communication [16, 20, 21]. The expression levels of Cx26 and Cx32, major connexins in the liver, are extremely low in several HCC lines, but Cx43, a minor connexin in the liver, is highly expressed in metastatic cancer [17, 20-22]. Cx43 knockdown using siRNA reduced cell proliferation and significantly suppressed the expression of Cx26 and Cox-2 in H69 cells stimulated with NDMA and ESP compare to silencing negative control stimulated with NDMA and ESP. The connexin proteins Cx43 and Cx26 are involved in cell modification upon stimulation by NDMA and ESP of *C. sinensis*.

In general, cells contain several known connexins, classified according to their intracellular location (Table 3).

**Table 3.**
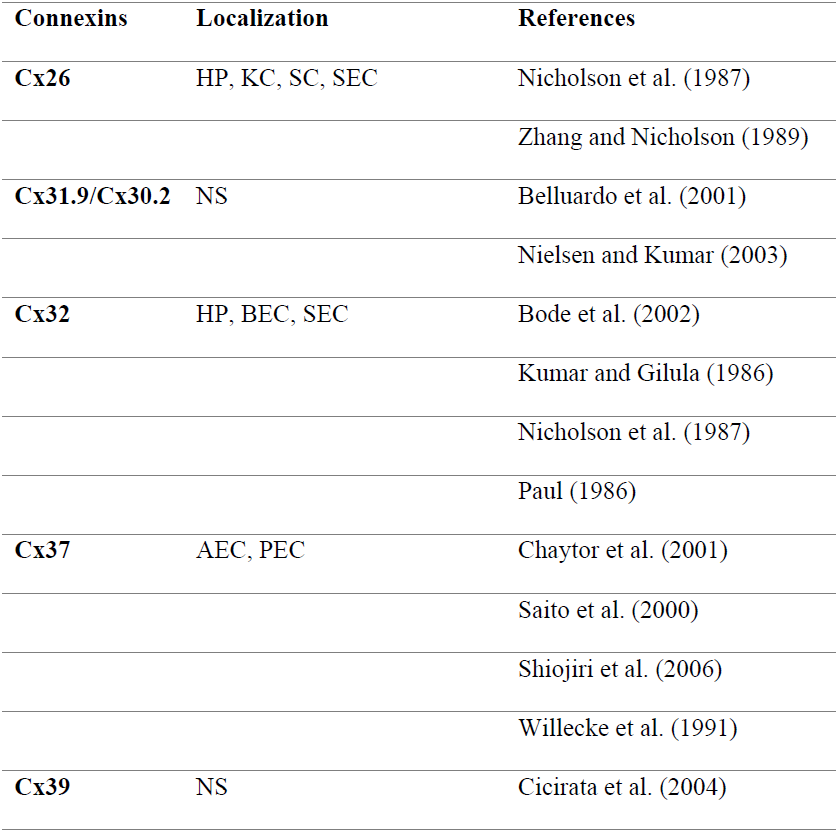
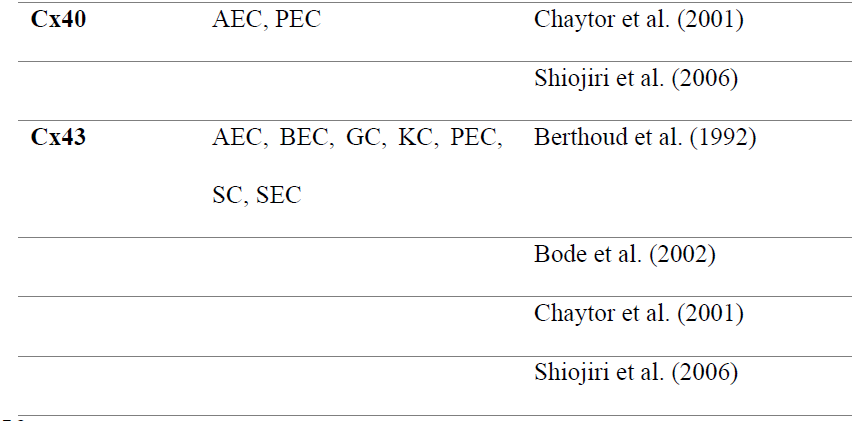
Full spectrum of connexins expressed in rodent and human livers

AEC, hepatic artery endothelial cell; BEC, biliary epithelial cell;

GC, Glisson’s capsule; HP, hepatocyte; KC, Kupffer cell; NS, not specified; PEC, portal vein endothelial cell; SC, stellate cell; SEC, sinusoidal endothelial cell. Intercellular communication via gap junctions is inhibited by increased Cox-2 expression, as is frequently observed in several forms of human malignancies [29, 30]. Recently, several reports have suggested that the carcinogenic mechanisms of hydrogen peroxide, TPA, and quinones may be involved in the inhibition of GJIC by Cx43 phosphorylation via ERK1/2 activation in rat liver epithelial cells [13, 17]. Furthermore, increased expression of Cx43 is positively correlated with NFκB activation in muscular arteries of patients undergoing coronary artery bypass graft surgery [31]. NFκB plays a central role in general inflammatory and immune responses. The 5’-flanking region of the *Cox-2* promoter contains an NFκB binding site, and NFκB is a critical regulator of Cox-2 expression in many cell lines [13, 31]. Taken together, the present findings suggest that Cx43 expression induces Cox2 over-expression via NFκB activation. However, until now, the link between NFκB activation, Cx43 expression, and Cox2 over-expression has not been clearly established. In the future, it will be interesting to examine the relationship between the GJIC and NFκB activation, Cox-2 by ESP of *C. sinensis* and NDMA.

In conclusion, our results suggest that Cx43 plays a key role in cell proliferation, potentially leading to CHCA development upon stimulation by ESP of *C. sinensis* and NDMA.

## Acknowledgments

We thank Professor Dae-Gon Kim of Chonbuk National University for providing H69 cells. We would like to thank Professor Sung-Jong Hong and Fuhong Dai, Department of Medical Environmental Biology, Chung-Ang University College of Medicine, for providing adult worms of *C*. *sinensis*.

